# Antiproliferative Activity of Cephalotaxus Esters: Overcoming Chemoresistance in Cancer Therapy

**DOI:** 10.1101/2023.11.06.565820

**Authors:** Vladimir Yong-Gonzalez, Constantine Radu, Paul A. Calder, David Shum, David Y. Gin, Mark G. Frattini, Hakim Djaballah

**Affiliations:** HTS Core Facility, Memorial Sloan Kettering Cancer Center, 1275 York Avenue, New York, NY 10065, USA; Molecular Pharmacology Program, Memorial Sloan Kettering Cancer Center, 1275 York Avenue, New York, NY 10065, USA; Department of Medicine, Memorial Sloan Kettering Cancer Center, 1275 York Avenue, New York, NY 10065, USA; Molecular Biology Program, Memorial Sloan Kettering Cancer Center, 1275 York Avenue, New York, NY 10065, USA; Deceased Memorial Sloan Kettering Cancer Center, 1275 York Avenue, New York, NY 10065, USA; Department of Analytical Chemistry, C16 Biosciences, New York, NY 10019, USA; Automation and Workflow Solutions, Molecular Devices, San Jose, CA 95134, USA; Department of Pediatrics, Memorial Sloan Kettering Cancer Center, New York, NY 10065, USA; Screening Discovery Platform, Institut Pasteur Korea, Seongnam-si, Gyeonggi-do, 13488, Republic of Korea; Office of the CMO, Cellectis Inc, New York, NY 10016, USA; Office of the CEO, Keren Therapeutics, New York, NY 10583, USA

**Keywords:** Homoharringtonine, multidrug resistance, leukemia, cancer, AML, CML

## Abstract

Omacetaxine, a semisynthetic form of Homoharringtonine (HHT), was approved for the treatment of chronic myeloid leukemia (CML). Previously, we published the synthesis of this natural alkaloid and three of its derivatives: deoxyharringtonine (DHT), deoxyhomoharringtonine (DHHT), and bis(demethyl)-deoxyharringtonine (BDHT); and reported on its refractory activity against the HL-60/RV+ cells over-expressing P-glycoprotein 1 (MDR1). In this study, we explored the extent of this resistance by first expanding the panel of established cell lines and second, using a panel of 21 leukemia patient derived primary cells. Here, we report a consistent resistance to HTT in K562 derived cells and in MES-SA/MX2 derived cells resistant to mitoxanthrone; all of them over-express MDR1, while we found U87MG-ABCG2 and H69AR cells to be very sensitive to HTT. In contrast, DHT, DHHT, and BDHT seemingly overcome this resistance due to the changes made to the acyl chain of HTT rendering the derivatives less susceptible to efflux. Surprisingly, the leukemia primary cells were very sensitive to HHT and its derivatives with low nanomolar potencies, followed by a new class of CDC7 kinase inhibitors, the anthracycline class of topoisomerase inhibitors, the DNA intercalator actinomycin-D, and the vinca alkaloid class of microtubule inhibitors. The mechanism of cell death induced by HTT and DHHT was found to be mediated via Caspase 3 cleavage leading to apoptosis. Taken together, our results confirm that HHT is a substrate for MDR1. It opens the door to a new opportunity to clinically evaluate HHT and its derivatives for the treatment of AML and other cancers.

## Introduction

Homoharringtonine (HHT) is a cephalotaxan alkaloid from *Cephalotaxus harringtonine*, an evergreen tree that grows in Southeast Asia.^1^ HHT forms a complex mixture with other less abundant cephalotaxan ester derivatives such as deoxyhomoharringtonine (DHHT), anhydroharringtonine (AHT) and deoxyharringtonine (DHT).^2^ The major molecular mechanism of action of HHT is the inhibition of protein synthesis,^3–5^ which is consistent with its property of binding to ribosome. It has been suggested that HHT inhibits the synthesis of short half-life proteins.^6^

Used for long time in traditional Chinese medicine, these alkaloids were first characterized in the 1970’s as part of the group of compounds that possess anti-neoplastic activity toward different types of leukemia cell lines.^7^ This discovery stimulated clinical trials to evaluate and validate their use in the treatment of leukemia.^8^ A phase II trial reported a complete remission for 28% of myeloid dysplasia syndrome (MDS) patients treated with HHT.^9^ Three independent studies on a single regimen of HHT and combination with interferon-α reported complete hematological remissions of 72, 92 and 85% of CML patients.^10–12^ HHT was also effective in the treatment of *de novo* AML with a report of complete remission of 83 % of the young adult subjected to this regime.^11^ Nevertheless, phase I and II studies found that HHT was less active in patients with relapsed AML or CML in blastic phase.^13–14^ AML patients with primary resistance to an initial anthracyclines/cytarabine combination did not respond at all to HHT treatment.^14^ Despite the lack of positive response for most patients, these studies reported a 16% complete remission for relapsed AML patients under HHT regimen and 23% for those treated under a combined regimen with cytarabine.^13–14^ Therefore, it seems that HHT efficacy will become reduced in both relapsed and refractory leukemia patients.

A majority of hematological malignancies show sensitivity to cytotoxic drugs during initial cycles of chemotherapy, but later develop multidrug resistance upon relapse.^15^ Studies on this mechanism of resistance identified multiple ABC transporters, such as multidrug resistance protein 1 (MRP1) and ATP-binding cassette super-family G member 2 (ABCG2), as major players of classical multidrug resistance (MDR).^16^ Importantly, the MDR1 transporter accounts for a frequency of about 30 to 50% of all AML cases refractory to treatments, making the need for novel clinical therapies imperative.^15^ Each transporter has multiple specific substrates and some of these substrates are shared with other transporters. For instance, MDR1 exhibits specificity toward paclitaxel and HHT,^17–18^ but is also active on the substrates of MRP1 and ABCG2, such as mitoxanthrone, doxorubicin, etoposide, and daunorubicin.^17^ Such a substrate redundancy is a common feature of ABC transporters and makes the discovery of new specific inhibitors a challenging process.

In an attempt to further explore the potential therapeutic effects of cephalotaxus ester derivatives, our group reported a novel synthetic approach to synthesize them using a novel convergent synthesis.^19^ These synthetic alkaloids proved to be cytotoxic against several tested cell lines from diverse origins including HL60/RV+, an MDR cell line derived from the human promyelocytic leukemia HL-60 cells.^19^ HL60/RV+ is known to over-express MDR1 and is resistant to a microtubule inhibitor vincristine. These cells were also 125-fold more resistant to HHT, a good substrate of MDR1, than HL-60, but sensitive to other cephalotaxus ester derivatives such as DHT which exhibited only 10-fold resistance more than HL-60. This rather low resistance index of DHT was also found to be the trend for others cephalotaxus ester derivatives such as DHHT and BDHT.^19^

These results prompted us to expand the established cell panel in order to investigate if DHHT, DHT and BDHT (**Fig. 1**) can overcome resistance for not being substrates of the multidrug efflux pumps, assess the chemosensitivity of a leukemia patient derived primary cells against a focused library of approved and experimental drugs including cephalotaxus esters, and investigate the mechanism of cell death caused by HTT and DHHT.

**Figure 1.**
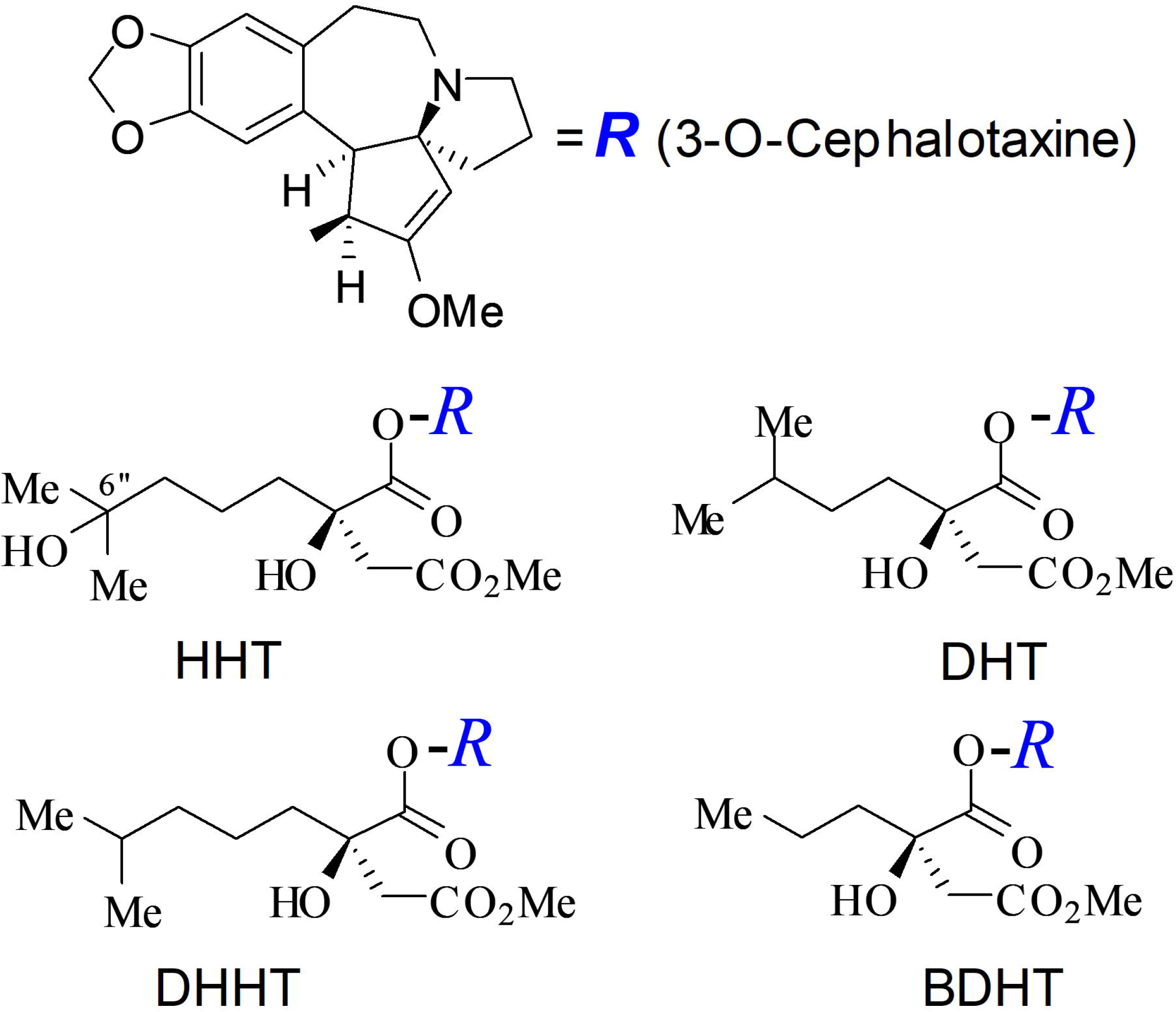
chemical structure of the cephalotaxus esters HHT, DHHT, DHT, and BDHT.

## Materials and Methods

### Reagents

All cell media to include Iscove’s Modified Dulbecco’s Medium (IMDM), RPMI 1640, Dulbecco’s modified Eagle’s medium (DMEM), L-Glutamine, Penicillin, Streptomycin, Phosphate Buffered Saline (PBS), and trypsin/EDTA were purchased from Life Technologies (Carlsbad, CA); normal goat serum, Hoechst 33342, and Alexa fluor 488 conjugated to streptavidin were purchased from Life Technologies. Heat-inactivated fetal bovine serum (FBS) was purchased from GE Healthcare (Chicago, IL). Bovine serum albumin (BSA) and Triton X-100 were purchased from Sigma-Aldrich (St. Louis, MO). The rabbit anti-cleaved Caspase 3 antibody was purchased from Cell Signaling Technology (Danvers, MA). Goat biotinylated anti-rabbit antibody was purchased from Vector laboratories (Newark, CA). Paraformaldehyde was obtained as a 32% (v/v) aqueous solution from Electron Microscopy Sciences (Hatfield, PA). All other reagents were available at the MSKCC HTS Core Facility or obtained as previously described.^19^

### Chemicals and drugs

The cephalotaxus ester derivatives, Homoharringtonine (HHT), deoxyhomoharringtonine (DHHT), deoxyharringtonine (DHT), and bis(demethyl)harringtonine (BDHT)) and hereafter named cephalotaxus ester derivatives were synthesize as previously described.^19^ All the other experimental and approved drugs (**Supp Table 1**) were obtained from the MSKCC HTS Core Facility, and subjected to a quality control step using mass spectrometry prior to their plating and use.

### Cell Lines used in this study

The following cell lines were purchased from ATCC (Gaithersburg, MD): K562 cells, HL-60 cells and its multidrug resistant derivations HL-60/MX1 and HL-60/MX2, human glioblastoma U87MG cells, human lung small cell carcinoma H69 cells and its multidrug resistant derivation H69AR, and human uterine sarcoma MES-SA cells and its multidrug resistant derivation MES-SA/MX2. U87MG-ABCG2 cells over-expressing the efflux pump ABCG2 were provided by Dr. Eric C. Holland (Fred Hutchinson Cancer Center, Seattle, USA). HL60/RV+ cells which are resistant to vincristine were provided by Dr. David A. Scheinberg (MSKCC, New York, USA). K-H200 and K-H400 cells were provided by Dr. Jean-Pierre Marie (University of Paris, France). These cells are MDR derivates of K562 cells and are resistant to HHT.^20^ The leukemia patient derived primary cells were isolated from patients at MSKCC and provided to us after obtaining patient written informed consent to the local Institutional Review Board approved protocol #06-107 for collection of tissue specimens. Cells were cultured either in IMDM, RPMI 1640, or DMEM, supplemented with 10% FBS, 100 units/mL penicillin, 100 μg/mL streptomycin, and 2mM L-glutamine. All cells were cultured under a humidified atmosphere at 37°C in an atmosphere containing 5% CO_2_.

### Dose response studies

Dose response of the compounds was assessed in duplicate as previously described.^19^ Twelve-point doubling dilutions of each compound in 10% DMSO (v/v) were prepared in an intermediate 384-well poly-propylene plate (Thermo Scientific, Waltham, MA) with 10 and 100 μM concentration as the upper limit. Five µL of each dilution was transferred to 384-well assay microtiter plates (Corning Inc., Somerville, MA) with a PP-384-MM Personal Pipettor equipped with a custom 384 head (Apricot Designs, Monrovia, CA). Cells were seeded in 45 µL suspensions with a Multidrop dispenser (Thermo Scientific, Waltham, MA) to each well which contained 5 μL compound dilution. The final concentrations of each compound were 12-point doubling dilutions with 1 and 10 μM as the highest concentration in 1% DMSO (v/v). Cell seeding densities used were as follows: 500cells/well for K562, U87MG and U87MG-ABCG2 cells; 1,000 cells/well for HL-60/MX1, HL-60/MX2, HL-60/RV+, K-H200, K-H400, HeLa-Empty, HeLa-Bcl-XL, MES-SA, MES-SA/MX2, H69, and H69AR cells; 2,000 cells/well for HL-60 cells. The patient derived primary leukemia cells were seeded at 10,000 cells/well. After an incubation of 72 hours (48 hours for leukemia primary cells) at 37°C, 5 µL of Alamar Blue was dispensed with a FlexDrop precision dispenser (Perkin Elmer, Boston, MA). After an additional incubation of 24 hours, fluorescence signal was measured on the LEADseeker™ Multimodality Imaging System (GE Healthcare, Piscataway, NJ).

Fluorescence signals in each well were compared to the average of the high and low controls which are located in the row A and P respectively of the same assay plate. The percentage of viability was calculated as follows: % Viability = 100 – (high control average – read value) / (high control average – low control average)*100. Dose response curves of triplicate sets were fitted by the logistic four-parameter equation of SigmaPlot (Systat Software Inc.). The average of three IC_50_ values represented the potency of each compound. Compound resistance of MDR cell lines relative to the parental cells was represented by Resistance index (RI).^19^

### Immunostaining of cleaved Caspase 3 in HeLa-Empty and HeLa-Bcl-XL cells

HeLa-Empty and HeLa-Bcl-XL cells^21^ (2.2 x 10^4^ cells) were seeded in 4-well glass chambered slide (Thermo Scientific, Waltham, MA) and incubated at 37°C. At 12 hours post cell seeding, the media was aspirated and fresh media containing 1 µM HHT or 1 µM DHHT was added and cells were incubated at room temperature for 8 hours. Cells were further incubated for 8 hours at 37 °C. The fresh medium used in each condition contains 1% DMSO (v/v) that was used as a carrier and a negative control. All of the incubation afterwards were conducted at room temperature except otherwise indicated. Cells were then fixed with 4% paraformaldehyde (v/v) in PBS for 20 minutes at room temperature and washed once with PBS.

The remaining experiments were conducted by the MSKCC Molecular Cytology Core Facility using the following steps: slides were treated for 15 minutes with 0.1% Triton X-100 (v/v) in PBS, and blocked with 10% normal goat serum and 2% BSA (w/v) in PBS for 30 minutes. The cells were incubated with a rabbit anti-cleaved Caspase 3 antibody (Cell Signaling Technology, Danvers, MA) at 4°C overnight in a moist chamber. After three washes with PBS, cells were incubated with goat biotinylated anti-rabbit antibody for 1 hour and the fluorescent reaction achieved by adding Alexa fluor 488 conjugated to streptavidin (Thermo Scientific, Waltham, MA) for 10 minutes. Subsequently, the slides were treated with 10 μM Hoechst solution in 0.05% (v/v) Triton X-100 in PBS for 15 minutes. After two washes with PBS, cells were kept at 4°C in the dark before imaging.

Image acquisition was acquired using a Leica TCS AOBS SP2 point-scanning confocal microscope (inverted stand) at 40X objective magnification using a Leica HC PL APO Oil Objective (40x/1.3). Images of nuclei stained with Hoechst in the blue channel were acquired using 405 nm excitation and 410-475 nm emission. Images of cleaved Caspase in the green channel were acquired using 488 nm excitation and 495-550 nm emission.

## Results

### Profiling cephalotaxus esters against sensitive and refractory HL-60 cells

We used the HL-60 cell line and its vincristine resistant HL-60/RV+ line and the two mitoxanthrone resistant cell lines, HL-60/MX1 and HL-60/MX2. We investigated their sensitivity to HHT, DHHT, DHT, BDHT, and 14 other FDA approved or experimental drugs (**Table 1**, **Fig. 2, Supp Table 2**). The potency of each compound against each cell line was presented as IC_50_ value and a compound resistance index (RI) of each cell line tested relative to the parental cell line as previously reported (19). As expected, the HL-60/RV+ cells confirmed their resistance to the microtubule inhibitor vincristine with an RI of 200 (**Supp Table 2**).^22^ This cell line also exhibited resistance toward other microtubule inhibitors like colchicine with an RI of 55 and vinblastine with an RI of 37. Similarly, it also showed resistance to the topoisomerase inhibitors daunorubicin with an RI of 52 and mitoxanthrone with an RI of 45 (**Supp Table 2**). We found that the HL-60/RV+ cells exhibited significant resistance toward HHT (RI = 37), but were very sensitive toward DHHT (RI= 5), DHT (RI= 5), and BDHT (RI = 4) (**Table 1**, **Fig. 2**).

**Figure 2.**
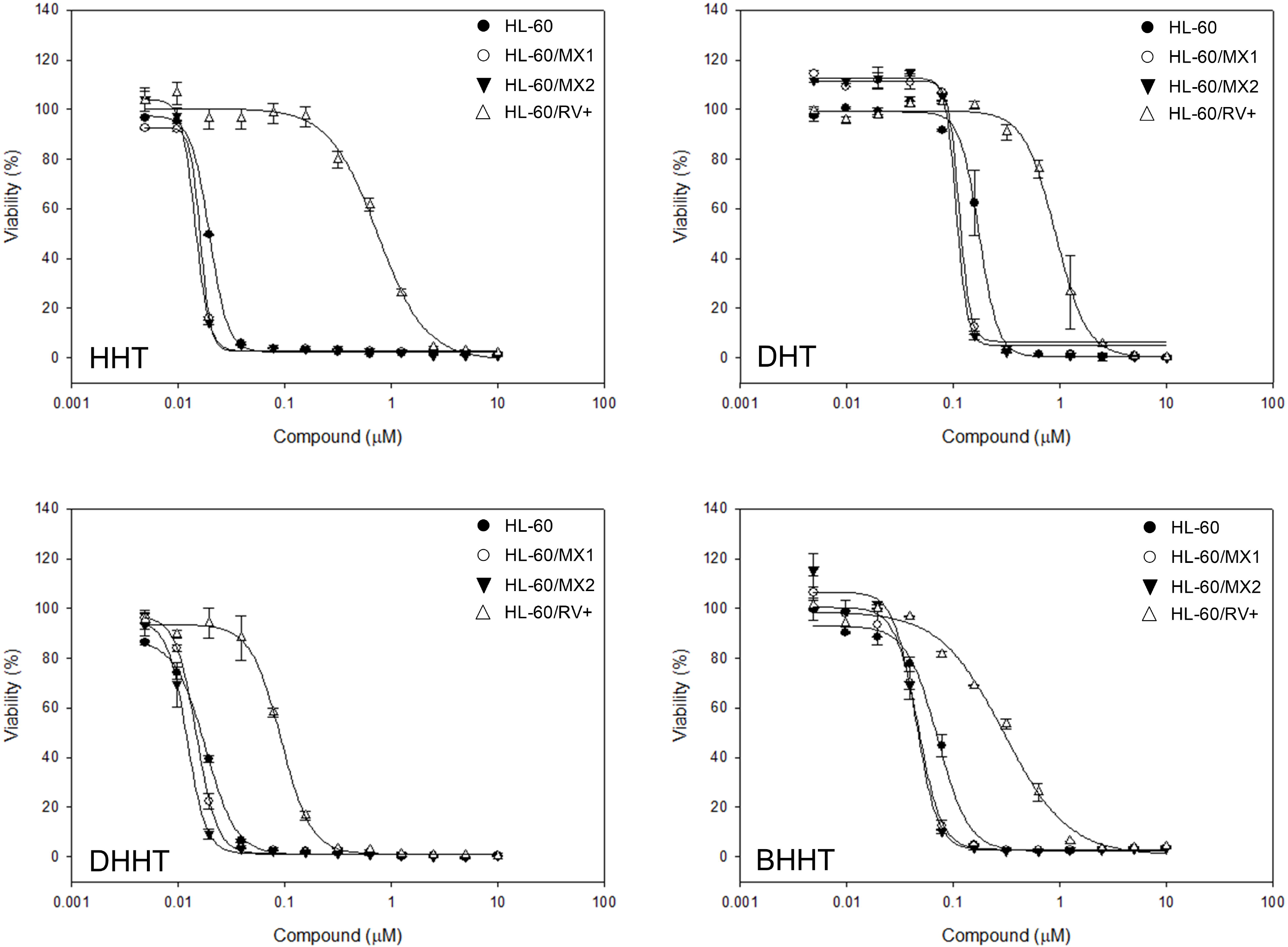
Comparative cytotoxic effects of cephalotaxus esters against sensitive (HL-60), Mitoxanthrone resistant (HL-60/MX1 and HL-60/MX2), and Vincristine resistant (HL-60/RV+) cells. Dose response studies of HHT, DHHT, DHT, and BDHT. The y-axis represents the percentage of cell viability that was calculated as described under Material and Methods.

**Table 1.**
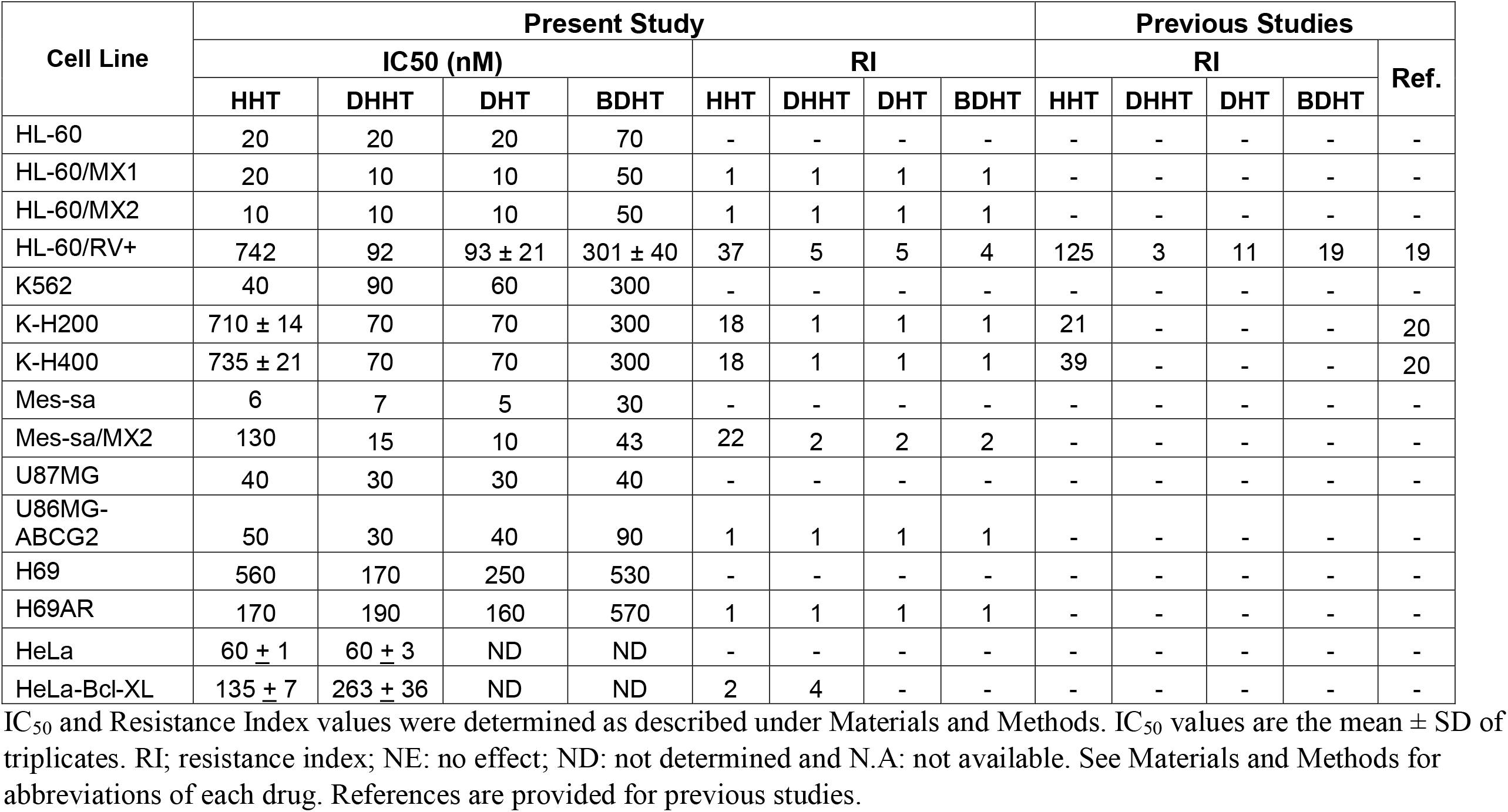
Cytotoxic activity and resistance profile of HHT, DHHT, DHT, and BDHT. IC_50_ and Resistance Index values were determined as described under Materials and Methods. IC_50_ values are the mean ± SD of triplicates. RI; resistance index; NE: no effect; ND: not determined and N.A: not available. See Materials and Methods for abbreviations of each drug. References are provided for previous studies.

In contrast to the HL-60/RV+ cells, the HL-60/MX1 and HL-60/MX2 cell lines exhibited a similar inhibitory activity profiles against HHT, DHT, DHHT, and BDHT as did the parental cell line HL-60 (**Table 1**, **Fig. 2**); and with no observed resistance (**Table 1**). They were found to be sensitive to mitoxanthrone with RIs of 12 and 10, respectively (**Supp Table 2**); it is expected as their reported resistance is mediated by reduction of expression and activity of the topoisomerase II enzyme, not by the over-expression of the MDR1 transporter.^23^ This result validates the observations that HHT resistance is mediated by the MDR1 transporter with an RI of 37 against the HL-60/RV+ cells known to over-express the efflux transporter MDR1.

### Profiling cephalotaxus esters against sensitive and refractory K562 cells

We next investigated the sensitivities of K562 cell line and its resistant lines, K-H200 and K-H400, which were established through viability selection with increasing concentrations of HHT,^20^ against a panel of selected FDA approved and experimental drugs including HHT, DHT, DHHT, and BDHT. As expected, the K-H200 and K-H400 cells were equally resistant to HHT with a RI of 18 as compared of the parental K562 cell line (**Table 1**; **Fig. 3; Supp Table 3**), in agreement with previously reported RI values of 21 and 39, respectively.^20^ Surprisingly, the K-H200 and K-H400 cells were found to be very sensitive to DHT, DHHT, and BDHT with IC_50_ values of 70, 70, and 300 nM, respectively; and similar to those obtained for the parental K562 cell line (**Table 1**; **Fig. 3**).

**Figure 3.**
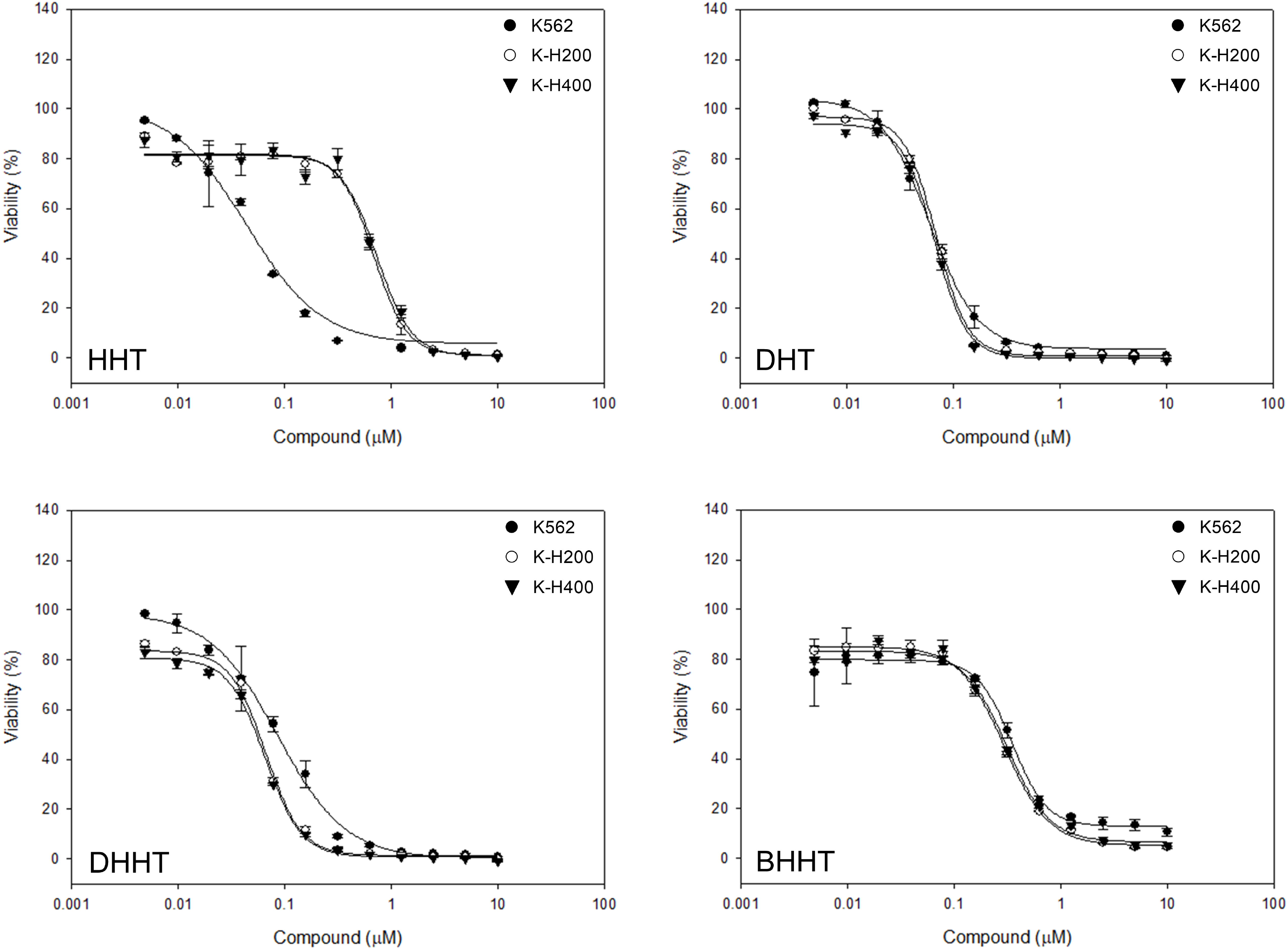
Comparative cytotoxic effects of cephalotaxus esters against sensitive (K562) and HHT resistant (K-H200 and K-H400) cells. Dose response studies of HHT, DHHT, DHT, and BDHT. The y-axis represents the percentage of viabilities that was calculated as described under Material and Methods.

K-H200 cells were found to be refractory to actinomycin-D, colchicine, vinblastine, vincristine, daunorubicin, doxorubicin, mitoxanthrone, and teniposide with RI values of 15, 108, 38, 158, 14, 16, > 41, > 28, respectively; while K-H400 cells were also found to be resistant with RI values of 14, 126, 35, 155, 18, 13, > 41, > 28, respectively (**Supp Table 3**). Zhou and co-workers reported RI values for vincristine, daunorubicin, mitoxanthrone, and etoposide of 508, 30, 24, and 4, respectively for K-H200 cells; and 857, 161, 53, and 16, respectively for K-H400 cells. ^20^ We noted that our RI values were not all in agreement with the published ones,^20^ but the trend of resistance for vincristine, daunorubicin, and mitoxanthrone was maintained. In our studies, etoposide was inactive, against both K-H200 and K-H400 cells, at the highest drug concentration tested of 10,000 nM (**Supp Table 3**).

The described mechanism of resistance by the K-H200 and K-H400 cells is via over-expression of the multidrug resistance gene MDR1, translating into a membrane bound p-glycoprotein functioning as an energy dependent efflux pump.^20^ Beside the four drugs reported by Zhou and Co-workers, we further identify actinomycin-D, colchicine, vinblastine, doxorubicin, and teniposide as substrates of the MDR1 efflux pump, and support the concept for the non-selectivity of the pump as these drugs represent different chemical classes (**Supp Table 3**); suggesting that resistance to HHT is mediated via the MDR1 glycoprotein.

### Profiling cephalotaxus esters against sensitive and refractory MES-SA cells

We investigated the sensitivities of the human uterine sarcoma cell line MES-SA and its mitoxanthrone resistant line MES-SA/MX2, against HHT, DHT, DHHT, and BDHT (**Table 1**). We found MES-SA cells to be very sensitive to HHT, DHT, DHHT, and BDHT with IC_50_ values of 6, 7, 5, and 30 nM, respectively; in contrast, the MES-SA/MX2 cells were much more resistant to the same compounds with IC_50_ values of 130, 15, 10, and 43 nM, respectively, and making it highly refractory to HHT with an RI value of 22 (**Table 1**). MES-SA/MX2 is a mitoxanthrone-resistant derivative of the human uterine sarcoma cell line MES-SA that displays features of overexpression of the two classical multidrug resistance proteins.^24^ This result suggests that HHT resistance is mediated via the MDR1 efflux pump.

### Profiling cephalotaxus esters against U87MG and U87MG-ABCG2 cells

We investigated the sensitivities of the human glioblastoma cell line U87MG and a derived line expressing the ABCG2 transporter U87MG-ABCG2,^25–26^ against HHT, DHT, DHHT, and BDHT (**Table 1**). We found U87MG cells to be sensitive to HHT, DHT, DHHT, and BDHT with IC_50_ values of 40, 30, 40, and 90 nM, respectively; similarly, the U87MG-ABCG2 cells were found to as sensitive to the same compounds with IC_50_ values of 50, 30, 40, and 90 nM, respectively. No resistance to HHT was observed with an RI value of 1 (**Table 1**). These data suggest that HHT is not a substrate for the ABCG2 transporter.

### Profiling cephalotaxus esters against H69 and H69AR cells

We investigated the sensitivities of the human small cell lung cancer cell line H69 and a its multidrug resistant variant H69AR obtained by culturing the H69 cells in gradually increasing doses of adriamycin up to 0.8 μM,^27^ against HHT, DHT, DHHT, and BDHT (**Table 1**). We found H69 cells to be relatively sensitive to HHT, DHT, DHHT, and BDHT with IC_50_ values of 560, 170, 250, and 530 nM, respectively; similarly, the multidrug resistant H69AR cells were found to be more sensitive to the same compounds with IC_50_ values of 170, 190, 160, and 570 nM, respectively. No resistance to HHT was observed with an RI value of 1, rather an enhanced chemo-sensitivity to HHT by H69AR cells (**Table 1**); unlike most other multidrug resistant cell lines, H69AR does not appear to express enhanced levels of P-glycoprotein, including MDR1, rather a reduced level of expression of topoisomerase II.^28^

### Profiling cephalotaxus esters against a panel of patient derived primary leukemia cells

In addition to the already established and well studied cell lines, we profiled a panel of 21 leukemia primary cells in different clinical stages of the disease, and obtained from 11 patients as some are serial cells from the same patient collected over time after relapsing from previous administered treatment regimens, against an expanded panel of experimental and FDA approved drugs (**Table 2**). It is of note that the cells were obtained from patients who are undergoing standard of care treatment or had undergone such treatment and none were naïve to chemotherapy. Remarkably, all of the cephalotaxus ester derivatives exhibited very potent cytotoxic activity, with IC_50_ values ranging from 6 to 145 nM (**Table 2**). Samples obtained from patient 1 were refractory to HHT, DHT, & DHHT, but very sensitive in the serial sample 1.3; Samples from patient 2 were refractory to DHT, but became sensitive in the serial sample 2.3. Similarly, patient 4 samples were refractory to BDHT, but became sensitive in the serial sample 4.2; this suggest in part that none of these AML patient derived samples were sensitive to HHT without indication of rapid resistance via the drug efflux pathways, and taking into account the small size of the cell panel.

**Table 2.** Efficacy of cephalotaxus ester derivatives against a panel of leukemia patient derived primary cells. IC_50_ values were determined as described under Materials and Methods, and are represented as mean ± SD of triplicates.

| Drug Name | Mode of Action | Patient Derived Leukemia Cells, IC50 (mM) |  |  |  |  |  |  |  |  |  |  |  |  |  |  |  |  |  |  |  |  |
| --- | --- | --- | --- | --- | --- | --- | --- | --- | --- | --- | --- | --- | --- | --- | --- | --- | --- | --- | --- | --- | --- | --- |
|  |  | 1.1 | 1.3 | 2.2 | 2.3 | 3.1 | 3.2 | 4.1 | 4.2 | 5.1 | 5.2 | 6.1 | 6.2 | 7.1 | 7.2 | 8.1 | 8.2 | 9.1 | 9.2 | 9.3 | 10.1 | 11.3 |
| HHT | Protein synthesis | 10 | 0.17 | 0.04 | 0.02 | 0.04 | 0.12 | 0.25 | 0.03 | 0.03 | 0.02 | 0.03 | 0.03 | 0.02 | 0.01 | 0.04 | 0.02 | 0.04 | 0.04 | 0.01 | 0.05 | 0.04 |
| DHT |  | 10 | 0.15 | 10 | 0.01 | 0.02 | 0.04 | 0.07 | 0.02 | 0.03 | 0.01 | 0.02 | 0.02 | 0.01 | 0.01 | 0.02 | 0.02 | 0.03 | 0.03 | 0.01 | 0.03 | 0.02 |
| DHHT |  | 10 | 0.13 | 0.02 | 0.02 | 0.02 | 0.04 | 0.07 | 0.02 | 0.02 | 0.02 | 0.03 | 0.02 | 0.01 | 0.01 | 0.06 | 0.03 | 0.04 | 0.03 | 0.02 | 0.04 | 0.04 |
| BDHT |  | 0.26 | 0.20 | 0.22 | 0.08 | 0.08 | 0.14 | 10 | 0.11 | 0.10 | 0.05 | 0.09 | 0.08 | 0.04 | 0.03 | 0.10 | 0.11 | 0.11 | 0.12 | 0.08 | 0.16 | 0.09 |
| Cytarabine | Anti-metabolite | 10 | 10 | 10 | 10 | 10 | 10 | 10 | 10 | 10 | 10 | 10 | 10 | 10 | 10 | 10 | 10 | 10 | 0.92 | 10 | 10 | 10 |
| 5-Fluorouracil |  | 10 | 10 | 10 | 10 | 10 | 10 | 10 | 10 | 10 | 10 | 10 | 10 | 10 | 10 | 10 | 10 | 10 | 10 | 10 | 10 | 10 |
| 6-Mercaptopurine |  | 10 | 10 | 10 | 10 | 10 | 10 | 10 | 10 | 10 | 10 | 10 | 10 | 10 | 10 | 10 | 10 | 10 | 10 | 10 | 10 | 10 |
| Methotrexate |  | 10 | 10 | 10 | 10 | 10 | 10 | 10 | 10 | 10 | 10 | 10 | 10 | 10 | 10 | 10 | 10 | 10 | 10 | 10 | 10 | 10 |
| Acivicin |  | 10 | 10 | 10 | 10 | 10 | 10 | 10 | 10 | 10 | 10 | 10 | 10 | 10 | 10 | 10 | 10 | 10 | 10 | 10 | 10 | 10 |
| Daunorubicin | DNA Topo inhibitor | 0.09 | 0.04 | 0.24 | 0.17 | 10 | 10 | 10 | 0.12 | 0.32 | 0.34 | 0.36 | 0.33 | 0.17 | 0.21 | 0.64 | 0.17 | 0.26 | 0.31 | 0.09 | 0.39 | 0.19 |
| Doxorubicin |  | 0.33 | 0.25 | 0.61 | 0.24 | 10 | 10 | 10 | 0.46 | 0.60 | 0.53 | 0.84 | 0.77 | 0.71 | 0.94 | 10 | 0.58 | 0.66 | 0.52 | 0.36 | 0.71 | 0.41 |
| Mitoxantrone |  | 0.16 | 0.09 | 0.29 | 0.29 | 10 | 10 | 10 | 0.15 | 0.49 | 0.62 | 0.51 | 0.45 | 0.16 | 0.12 | 10 | 0.19 | 0.40 | 0.19 | 0.16 | 0.45 | 0.35 |
| Camptothecin |  | 10 | 10 | 10 | 10 | 0.02 | 10 | 10 | 10 | 10 | 10 | 10 | 10 | 10 | 10 | 10 | 10 | 10 | 10 | 10 | 10 | 10 |
| Teniposide |  | 10 | 0.69 | 10 | 10 | 10 | 10 | 10 | 10 | 10 | 10 | 10 | 10 | 10 | 10 | 10 | 10 | 10 | 10 | 10 | 10 | 10 |
| Etoposide |  | 10 | 10 | 10 | 10 | 10 | 10 | 10 | 10 | 10 | 10 | 10 | 10 | 10 | 10 | 10 | 10 | 10 | 10 | 10 | 10 | 10 |
| Actinomycin-D | DNA intercalating agent | 0.13 | 10 | 0.13 | 0.05 | 10 | 10 | 0.42 | 10 | 0.09 | 10 | 0.14 | 0.13 | 0.18 | 0.16 | 0.30 | 0.24 | 0.11 | 0.10 | 0.18 | 0.15 | 0.10 |
| m-AMSA |  | 0.57 | 0.39 | 1.22 | 1.51 | 10 | 10 | 10 | 10 | 10 | 10 | 10 | 10 | 10 | 10 | 10 | 10 | 10 | 10 | 10 | 10 | 10 |
| Carboplatin | DNA alkylating agent | 10 | 10 | 10 | 10 | 10 | 10 | 10 | 10 | 10 | 10 | 10 | 10 | 10 | 10 | 10 | 10 | 10 | 10 | 10 | 10 | 10 |
| Melphalan |  | 10 | 10 | 10 | 10 | 10 | 10 | 10 | 10 | 10 | 10 | 10 | 10 | 10 | 10 | 10 | 10 | 10 | 10 | 10 | 10 | 10 |
| Vinblastine | Microtubule inhibitor | 0.10 | 0.11 | 0.48 | 0.13 | 10 | 10 | 10 | 10 | 10 | 10 | 0.15 | 10 | 0.14 | 0.06 | 10 | 10 | 10 | 0.19 | 0.14 | 0.09 | 10 |
| Vincristine |  | 0.12 | 0.09 | 10 | 0.19 | 10 | 10 | 10 | 10 | 10 | 10 | 0.22 | 0.23 | 0.02 | 0.02 | 0.22 | 0.20 | 0.29 | 0.20 | 0.19 | 10 | 10 |
| Colchicine |  | 0.05 | 0.04 | 0.13 | 0.13 | 10 | 10 | 10 | 0.09 | 10 | 10 | 0.12 | 10 | 0.09 | 0.05 | 10 | 0.07 | 10 | 10 | 10 | 0.06 | 10 |
| Paclitaxel |  | 10 | 10 | 10 | 10 | 10 | 10 | 10 | 10 | 10 | 10 | 10 | 10 | 10 | 10 | 10 | 10 | 10 | 10 | 10 | 10 | 10 |
| MSK-747 | CDC7 kinase inhibitor | 0.21 | 0.21 | 0.08 | 0.06 | 0.26 | 0.34 | 0.23 | 0.49 | 0.40 | 0.19 | 0.37 | 0.36 | 0.07 | 0.14 | 0.78 | 0.69 | 0.56 | 0.49 | 0.67 | 0.21 | 0.20 |
| MSK-777 |  | 0.13 | 0.13 | 0.03 | 0.03 | 0.12 | 0.19 | 0.17 | 0.20 | 0.31 | 0.11 | 0.30 | 0.27 | 0.03 | 0.04 | 0.40 | 0.39 | 0.41 | 0.34 | 10 | 0.15 | 0.13 |

The same primary cell panel was also very sensitive to two experimental inhibitors, MSK-747 and its derivative MSK-777, of the cell division cycle 7 (CDC7), serine threonine kinase, that is critical in the regulation of cell cycle progression, and with activities in the nanomolar range., Only serial sample 9.3 did not respond to MSK-777, though there were sensitive in the earlier 9.1 and 9.2 samples, while remaining sensitive to MSK-747 (**Table 2**). This result validates the critical importance of CDC7 in the survival of tumor cells and makes this a very promising target for clinical development.

In complete contrast, most of the cells in the primary leukemia cell panel were not sensitive to anti-metabolite drugs, including cytarabine and 5-fluorouracil, DNA alkylating agents including carboplatin and melphalan; and some topoisomerase inhibitors including campothecin, teniposide, and etoposide (**Table 2**). Interestingly, etoposide and teniposide are both synthetic and less toxic derivatives of podophyllotoxin; whereas camptothecin, which remains an experimental drug. The taxane microtubule inhibitor paclitaxel showed no cytotoxic activity against the panel of 21 patient leukemia primary cells, whereas the vinca alkaloid based inhibitors, vincristine and vinblastine, showed some potent activity but not across all the primary cells. Moreover, colchicine, an alkaloid isolated from *Colchicum autumnale*, exhibited similar activity to both vincristine and vinblastine (**Table 2**).

The anthracycline family of topoisomerase inhibitors tested, including daunorubicin, doxorubicin and mitoxanthrone, showed very potent activities with IC_50_ values ranging from 130 to 950 nM, except for primary leukemia cell samples from patient 3 that were totally refractory to the three drugs, while samples from patients 4 and 8 were refractory to doxorubicin and mitoxanthrone but became sensitive in the obtained serial sample 4.2 and 8.2 (**Table 2**), making these patient derived cells were more sensitive to anthracyclines, followed by podophyllotoxins and by camptothecins.

### Cephalotaxus esters induce apoptosis via Caspase 3 activation

To investigate the mechanism by which HHT & DHHT mediate their cytotoxic activity, we used the isogenic pair of human cervical tumor cells HeLa-Bcl-XL and HeLa-Empty. The HeLa-Bcl-XL cells stably over-express the anti-apoptotic protein Bcl-XL from an expression plasmid, and HeLa-Empty cells stably maintain an empty control vector.^21^ These cells are suitable cells to elucidate the mechanism of cytotoxicity mediated by HHT & DHHT as they allow the comparison of the levels of cleaved Caspase 3 signal in the presence or absence of Bcl-XL over-expression. HeLa cells were found to be very sensitive to both HHT and DHHT with IC_50_ values of 60 ± 1 nM and 60 ± 3 nM, respectively; while HeLa-Bcl-XL cells appear to be less sensitive with IC_50_ values of 135 ± 7 nM and 265 ± 36 nM, respectively (**Table 1**).

Cells were treated for 8 hours with 1% DMSO (v/v) as a control, 1 µM HHT, or 1 µM DHHT, after which, cells were fixed and stained with an anti cleaved Caspase 3 antibody and imaged (**Fig. 4**). Minimally Caspase 3 cleavage was detected in the group treated with DMSO for HeLa-Empty & HeLa-Bcl-XL (**Fig. 4**), and it was clearly attenuated in the treated HeLa-Bcl-XL cells (**Fig. 4**), confirming that the mechanism of cell death, upon treatment, is mediated via apoptosis. The Caspase 3 cleavage intensities were also very high in both the HHT and the DHHT treated group for HeLa-Empty, while attenuated in the HeLa-Bcl-XL treated group of cells (**Fig. 4**)

**Figure 4.**
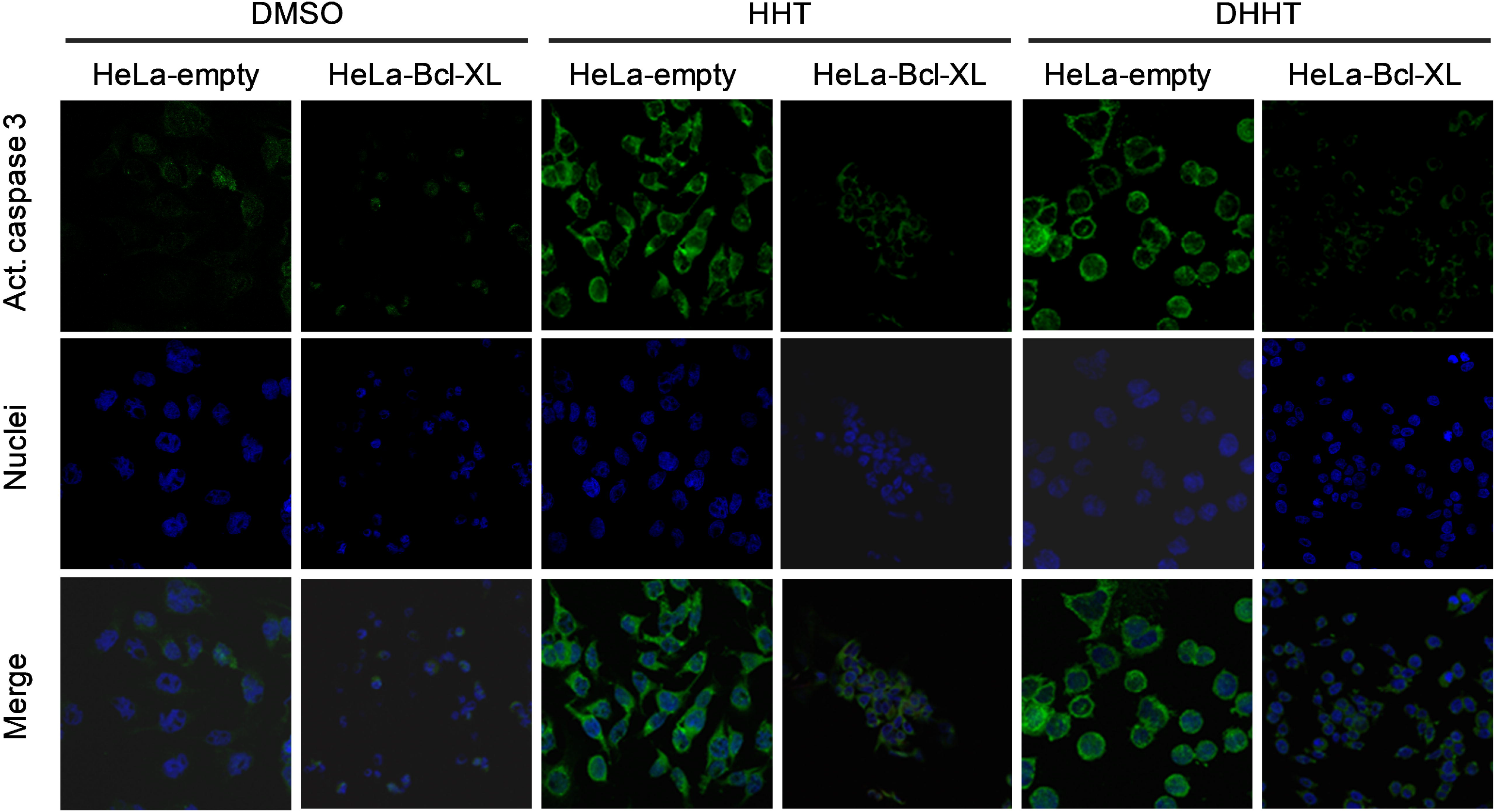
HHT and DHHT induce Caspase 3 cleavages in HeLa cells. The isogenic pair HeLa-Empty and HeLa-Bcl-XL cells were treated for 8 hours at 37 °C with: 1% DMSO (v/v), 1 µM HHT, or 1 µM DHHT. After fixation, cleaved Caspase 3 and nuclei were stained with anti-cleaved Caspase 3 antibody (green channel) and Hoechst 33342 (blue channel), respectively, and imaged with a 40X oil magnifying objectives. Top row represents the green channel, middle row, the blue channel, and the bottom row, the overlay of both channels.

In the HeLa-Empty cells, high intensity of the green fluorescence associated to the cleavage of Caspase 3 was observed, showing that these compounds work through a mechanism that activates apoptosis (**Fig. 4**). In contrast, this green fluorescent signal is significantly attenuated in HeLa-Bcl-XL cells. Nonetheless, these cells still showed higher levels of Caspase 3 cleavage when compared to the negative control treated with 1% DMSO (v/v) (**Fig. 4**). Possibly, the expression of Bcl-XL is not sufficient to suppress the cleavage of Caspase 3 in the HeLa-Bcl-XL cells.

## Discussion

HHT is a naturally occurring alkaloid originally isolated from the cephalotaxus hainanensis.^1^ Omacetaxine, a semisynthetic form of HHT, has been approved by the US Food and Drug Administration (FDA) for treatment of CML with resistance and/or intolerance to imatinib or other tyrosine kinase inhibitors.^29^ However, it is still awaiting approval for use in the treatment of AML. The elucidation of the function of P-glycoprotein has had a major impact on the understanding of multidrug resistance in cancer. More importantly, it has allowed the investigational merits of rational strategies aimed at reversing the emerging resistance in the clinic.^30^ Nevertheless, it has become clear that overexpression of P-glycoprotein is likely to be only one of several resistance mechanisms.^31^

With that in mind, we have investigated existing and potential chemo-resistance of the cephalotaxus ester HHT and its derivatives DHT, DHHT, and BDHT utilizing, first, an already established and characterized panel of cell lines, including cells with resistance to Mitoxanthrone, HL-60/MX1, HL-60/MX2, and MES-SA/MX2, resistance to vincristine, HL-60/RV+, resistance to HHT, K-H200 and K-H400, resistance to adriamycin, H69AR, and over-expression of the ABCG2 transporter, U87M-ABCG2 (**Table 1**); and second, a panel of twenty one patient derived primary AML cells were assessed for their sensitivities (**Table 2**).

We confirm prior findings that HTT is a substrate for MDR1,^19^ causing its reduced cytotoxic effects against HL-60/RV+, K-H200, K-H400, and MES-SA/MX2 cells; and with RI values of 37, 18, 18, and 22, respectively. No effect of the cytotoxic activity of HTT was observed when the U86MG-ABCG2 cells were used, suggesting that HHT is not a substrate for the ABCG2 efflux pump; more so that similar observations were made with the adriamycin resistant cell line H69AR, potentially ruling out the topoisomerase II activity as means of detoxifying the H69 cells from the cytotoxic effects of HHT. Combined, these data suggest that potential liabilities for HHT resistance will most likely due to expression of MDR1. It is of note that the MDR1 transporter accounts for up to 50% of all refractory AML cases, making it a priority for future clinical investigations.^15^

The chemo-sensitivity profile of DHT, DHHT, and BDHT is more favorable as the refractory cells remain more sensitive with RI values ranging from 1 to 5 (**Table 1**), even for the K-H200 and K-H400 cells that were obtained by resistance selectivity to HTT.^20^ The differential cytotoxic activity between HHT and its derivatives is consistent with the structural differences in their acyl chain (**Fig. 1**). In fact, variations in the acyl chain structure account for the increased hydrophobicity of DHHT, DHT and BDHT making them less susceptible to the activity of the MDR1 transporter.^19^ Therefore, modification of the acyl chain of the cephalotaxus esters is a valid strategy to generate new derivatives with reduced liability for MDR1 efflux.

We previously reported on the successful development of a high throughput assay of cellular proliferation and its validation through screening established immortalized cells as models for leukemogenesis and lymphomagenesis, against a 2,800 compound library containing nearly all of the currently used chemotherapeutics in addition to some potential novel agents.^32^ Differential compound responses were observed across the tested cell types indicating that they can be clustered by their sensitivities to small molecules with conserved chemical scaffolds, allowing for the platform usage to broaden our understanding of these molecular signatures by screening primary patient-derived leukemia cells harboring identifiable molecular lesions, with discovering novel chemical scaffolds for development as therapeutics for leukemias.^32^

In this study, we performed a comprehensive screen of a panel of 21 patient derived primary leukemia cells obtained from eleven patients against a focused compound set of 18 approved and 7 experimental drugs, to include HHT derivatives DHT, DHHT, and BDHT; and the CDC7 kinase inhibitors MSK-747 and MSK-777 (**Supp Table 1**). Differential chemosensitivities were observed with no cytotoxic activity for the anti-metabolites group including cytarabine, the DNA alkylating agents, carboplatin and melphalan, some of the DNA topoisomerase inhibitors such as camptothecins, teniposide, and etoposide, and the microtubule inhibitor paclitaxel (**Table 2**). In contrast, the anthracycline chemical class of topoisomerase inhibitors, namely, daunorubicin, doxorubicin, and mitoxanthrone showed good activity with the exception of patient 3 samples which were completely refractory, while patient sample 4.1 was refractory, but a later sample 4.2 became very sensitive to these drugs. The CDC7 kinase inhibitor MSK-747 and its pro-drug MSK-777 exhibited very good cytotoxic activity across the patient derived samples, with perhaps patient sample 9.3 which became refractory to MSK-777, while it was sensitive at earlier time points for sample 9.1 and 9.2 (**Table 2**). Interestingly, both MSK-747 and MSK-747 belong to the anthracycline class of molecules. ^33^

Overwhelming responses were observed for HHT and its derivatives, and with only partial responses for patient samples 1, 2, and 4, which became sensitive with time as shown by the latter samples 1.3, 2.3, and 4.2 (**Table 2**). This was an unexpected result for HHT as we predicted that at least 50% of the patient’s derived samples would have been refractory. A possible explanation could be that these primary leukemia cells do not express the MDR1 transporter that promotes sufficient resistance against most of the compounds present in the panel. In fact, expression of MRP1 and ABCG2 transporters have been detected in AML patient derived cells albeit its clinical relevance remains to be seen.^34^ Another possibility would correlate the emergence of resistance with a brief exposure of the patients to chemotherapeutic drugs in initial treatment cycles of chemotherapy.^35^ A rapid activation (within minutes) of MDR has been documented in patients with metastatic sarcoma after exposure to doxorubicin and a similar phenomenon was also observed in patients receiving the histone deacetylase inhibitor FR901228.^35–36^ However, this possibility is ruled out because it does not explain the remarkable sensitivity of the primary leukemia cells to HHT, which is a validated specific substrate of MDR1.^20,37^ Therefore, it is likely that these primary leukemia cells do not express MDR1 or its expression is not enough to mediate a significant resistance to HHT. Importantly, none of these hypotheses are relevant to explain the notably sensitivity of these cells toward the DHHT, DHT and BDHT as these compounds are not susceptible to MDR1, MRP1 (also expressed by K-H200) or ABCG2 (**Table 1**).

The cephalotaxus ester derivatives were the more potent compounds against the leukemia primary cells with nanomolar activity and sufficient to manifest their remarkable cytotoxicity effects. Clinical trials showed the maximal tolerated dose of HHT administered in AML patients generate a peak plasma concentration of 175 ± 37 nM. Therefore, the concentration of cephalotaxans derivatives including HHT achieved in our study is consistent with those reported in clinical trials.^38–39^

We investigated the mechanism of cytotoxicity of the cephalotaxus ester HHT and DHHT using HHT the HeLa-Empty and HeLA-Bcl-XL cell system. This system has been previously validated to verify that Mcl-1 degradation and Bax/Bcl-XL translocation to the mitochondria is a sequential event required for the final step of apoptosis.^21^ These cells (HeLa-Bcl-XL) do not release cytochrome c from the mitochondria, so cleavage of caspase 3 is suppressed by overexpression of Bcl-XL. In these experimental conditions, HHT and DHHT induced significant caspase 3 cleavage in the absence of Bcl-XL **(Fig. 4)**. This negative correlation between the levels of caspase 3 cleavages and the expression of the antiapoptotic protein Bcl-XL suggest that these compounds induce cytotoxicity through apoptosis. Most importantly, it demonstrated that DHHT parallels HHT in its ability to induce caspase 3 cleavages in these cells. This result correlates well with the fact that HHT is very susceptible to the activity of the MDR transporter.^17–18^ Taken together, these results corroborate that DHHT cytotoxicity involves the cleavage of Caspase 3 as means to induce and activate apoptosis as the mechanism of cell death.

In HeLa-Empty/HeLa-Bcl-XL cells, genotoxic stress induced by etoposide or gamma radiation causes protein synthesis inhibition. This inhibition leads to the depletion of the sensor Mcl-1 and triggers apoptosis.^21^ We speculate that similar mechanisms could be operating when the cells are exposed to DHHT or the other cephalotaxus ester derivatives. In support of this hypothesis, it has been shown that HHT induces apoptosis in HL-60 cells as well as primary cells derived from CML and AML patients.^39–40^ Importantly, the underlying mechanism is through the depletion of Mcl-1 protein.^39^ Depletion of Bcr-Abl by HHT treatment has been found to induce cytotoxicity in K562 cells.^40^

## Conclusion

We report on the expansion of our studies into the potential resistance of HHT cytotoxic activity via the MDR1 efflux pump and we show that though HHT is a good MDR1 substrate, its derivatives, DHT, DHT, and BDHT, were found to overcome this resistance due to the acyl chain modification (**Fig. 1**) rendering them less susceptible to efflux. We report that Caspase 3 activation was initiated upon treatment with HHT and DHHT and leading to cell death as the mechanism of cell death. We further report on the differential cytotoxic activity of a focused library of approved and experimental drugs against a panel of leukemia patient derived primary samples; notably, the very potent activity of all the cephalotaxus esters across various patient derived primary leukemia cells with the observation that if samples were refractory at one time point, they became sensitive at a follow up time point. Finally, HTT provides an excellent avenue for cancer therapeutics for AML, CML, and other cancers, and if ever resistance develops in patients receiving HHT, its derivatives can be considered as second generation cephalotaxus ester cancer therapeutic.

## Acknowledgements

We thank Drs Eric C. Holland (Fred Hutchinson Cancer Center, Seattle, USA), David A. Scheinberg (MSKCC, New York, USA), and Jean-Pierre Marie (University of Paris, France) for sharing valuable cells enabling our studies. We also thank members of the MSKCC Molecular Cytology Core Facility for their help with the confocal imaging experiments.

## Authors’ Contributions

Conceptualization: DYG, MGF, and HD; Methodology: VYG, MGF, and HD; Experimental Execution: VYG, JTM, CR, PAC, DYG, and HD; Data Analysis: VYG, MGF and HD; Manuscript Writing: VYG, MGF and HD; Supervision: HD.

