## Supplemental Table 1 for "Antiproliferative Activity of Cephalotaxus Esters: Overcoming Chemoresistance in Cancer Therapy"

**Supplementary Table 1** List of compounds used in profiling the leukemia patient derived samples.

| **Drug Name** | **Approval status** | **Mode of Action** |
| --- | --- | --- |
| Homoharringtonine, HHT | Approved | Protein synthesis |
| Deoxyharringtonine, DHT | Experimetal |  |
| Deoxyhomoharringtonine, DHHT | Experimetal |  |
| Bis(demethyl)deoxyharringtonine, BDHT | Experimetal |  |
| Cytarabine | Approved | Anti-metabolite |
| 5-Fluorouracil | Approved |  |
| 6-Mercaptopurine | Approved |  |
| Methotrexate | Approved |  |
| Acivicin | Experimental |  |
| Daunorubicin | Approved | DNA Topo inhibitor |
| Doxorubicin | Approved |  |
| Mitoxantrone | Approved |  |
| Camptothecin | Approved |  |
| Teniposide | Approved |  |
| Etoposide | Approved |  |
| Actinomycin-D | Approved | DNA intercelating agent |
| m-AMSA | Approved |  |
| Carboplatin | Approved | DNA alkylating agent |
| Melphalan | Approved |  |
| Vinblastine | Approved | Microtubule inhibitor |
| Vincristine | Approved |  |
| Colchicine | Experimental |  |
| Paclitaxel | Approved |  |
| MSK-747 | Experimental | CDC7 kinase inhibitor |
| MSK-777 | Experimental |  |
