## Supplemental Table 2 for "Antiproliferative Activity of Cephalotaxus Esters: Overcoming Chemoresistance in Cancer Therapy"

**Supplementary Table 2** Resistance profile of HL-60 derived cell lines HL-60/MX1, HL-60/MX2, and HL60/RV+.

| **Drug Name** | **Present Study** | | | | | | | **Previous Studies** | | | |
| --- | --- | --- | --- | --- | --- | --- | --- | --- | --- | --- | --- |
|  | **IC50 (nM)** | | | | **RI** | | | **RI** | | | |
|  | **HL-60** | **HL-60/MX1** | **HL-60/MX2** | **HL-60/RV+** | **HL-60/MX1** | **HL-60/MX2** | **HL-60/RV+** | **HL-60/MX1** | **HL-60/MX2** | **HL-60/RV+** | **Ref.** |
| Actinomycin-D | 40 | 100 | N.E. | 340 | 3 | N.D. | 9 | - | 2 | - | 23 |
| Camptothecin | 40 | 30 | 20 | N.E. | 1 | 0.5 | N.D. | - | - | - | - |
| Colchicine | 10 | 10 | 10 | 550 | 1 | 1 | 55 | - | - | - | - |
| Cytarabine | 210 | 50 | 50 | N.E. | 0.2 | 0.2 | N.D. | - | - | - | - |
| Daunorubicin | 30 | 100 ± 14 | 95 ± 7 | 1,570 | 3 | 3 | 52 | - | 4 | - | 23 |
| Doxorubicin | 215 ± 64 | 110 | 110 | 2,140 | 0.5 | 0.5 | 10 | 3 | 4 | - | 23 |
| Etoposide | 300 | 2,300 | 2,110 | N.E. | 7 | 7 | N.D. | 11 | 15 | - | 23 |
| m-AMSA | 125 ± 7 | 550 ± 120 | 640 ± 141 | N.E. | 4 | 5 | N.D. | 10 | 32 | - | 23 |
| Methotrexate | N.E. | N.E. | N.E. | N.E. | N.D. | N.D. | N.D. | - | - | - | 23 |
| Mitoxanthrone | 10 | 120 | 100 | 455 ± 78 | 12 | 10 | 45 | 10 | 35 | - | 23 |
| Paclitaxel | N.E. | 270 | 270 | N.E. | N.D. | N.D. | N.D. | - | - | - | - |
| Teniposide | 40 | 270 ± 14 | 270 ± 14 | N.E. | 7 | 7 | N.D. | - | 24 | - | 23 |
| Vinblastine | 10 | 10 | 10 | 370 | 1 | 1 | 37 | 1 | 1 | - | 23 |
| Vincristine | 10 | 10 | 10 | 2,015 ± 78 | 1 | 1 | 200 | 1 | 1 | - | 23 |

IC_50_ and Resistance Index values were determined as described under Materials and Methods. IC_50_ values are mean ± SD of triplicates. References are provided for previous studies. NE: no effect; ND: not determined.
