## Supplemental Table 3 for "Antiproliferative Activity of Cephalotaxus Esters: Overcoming Chemoresistance in Cancer Therapy"

**Supplementary Table 3** Resistance profile of K562 derived cell lines K-H200 and K-H400.

| **Drug Name** | **Present Study** | | | | | **Previous Study** | | |
| --- | --- | --- | --- | --- | --- | --- | --- | --- |
|  | **IC50 (nM)** | | | **RI** | | **RI** | | |
|  | **K562** | **K-H200** | **K-H400** | **K-H200** | **K-H400** | **K-H200** | **K-H400** | **Ref.** |
| Actinomycin-D | 40 | 615 ± 92 | 560 | 15 | 14 | - | - | - |
| Colchicine | 10 | 1,080 ± 42 | 1,255 ± 7 | 108 | 126 | - | - | - |
| Cytarabine | N.E. | N.E. | N.E. | N.D. | N.D. | - | - | - |
| Daunorubicin | 130 ± 14 | 1,825 ± 92 | 2,300 ± 71 | 14 | 18 | 30 | 161 | 20 |
| Doxorubicin | 135 ± 35 | 2,140 ± 212 | 1,740 ± 198 | 16 | 13 | - | - | - |
| Etoposide | N.E. | N.E. | N.E. | N.D. | N.D. | 4 | 16 | 20 |
| Methotraxate | N.E. | N.E. | N.E. | N.D. | N.D. | - | - | - |
| Mitoxanthrone | 240 ± 42 | > 10,000 | > 10,000 | > 41 | > 41 | 24 | 53 | 20 |
| Paclitaxel | N.E. | N.E. | N.E. | N.D. | N.D. | - | - | - |
| Teniposide | 360 ± 57 | > 10,000 | > 10,000 | > 28 | > 28 | - | - | - |
| Vinblastine | 10 | 375 ± 21 | 350 ± 14 | 38 | 35 | - | - | - |
| Vincristine | 10 | 1,575 ± 78 | 1,550 ± 198 | 158 | 155 | 508 | 857 | 20 |

IC_50_ and Resistance Index values were determined as described under Materials and Methods. IC_50_ values are mean ± SD of triplicates. References are provided for previous studies. NE: no effect; ND: not determined.
